# Expression of progerin does not result in an increased mutation rate

**DOI:** 10.1101/047506

**Authors:** Emmanuelle Deniaud, Shelagh Boyle, Wendy Bickmore

## Abstract

In the premature ageing disease Hutchinson-Gilford progeria syndrome (HGPS) the underlying genetic defect in the *lamin A* gene leads to accumulation at the nuclear lamina of progerin – a mutant form of lamin A that cannot be correctly processed. This has been reported to result in defects in the DNA damage response and in DNA repair, leading to the hypothesis that, as in normal ageing and in other progeroid syndromes caused by mutation of genes of the DNA repair and DNA damage response pathways, increased DNA damage may be responsible for the premature ageing phenotypes in HGPS patients. However, this hypothesis is based upon the study of markers of the DNA damage response, rather than measurement of DNA damage per se or the consequences of unrepaired DNA damage -mutation. Here, using a mutation reporter cell line, we directly compared the inherent and induced mutation rates in cells expressing wild-type lamin A or progerin. We find no evidence for an elevated mutation rate in progerin-expressing cells. We conclude that the cellular defect in HGPS cells does not lie in the repair of DNA damage per se.

## Introduction

Hutchinson-Gilford progeria syndrome (HGPS) is a dominant severe premature ageing genetic disease characterised by the appearance in childhood of age-related symptoms such as hair loss, thin skin, vascular defects and atherosclerosis. In the majority of cases, the underlying genetic defect is a point mutation that results in activation of a cryptic splice site in exon 11 of *lamin A* (*LMNA*) (1). This leads to accumulation of lamin A protein lacking the 50 amino acid (a.a.) domain toward the C-terminus (LAΔ50/progerin) that is required for the final endoproteolytic cleavage step during the normal processing of pre-lamin A to lamin A. As a consequence, LAΔ50 remains permanently farnesylated (2,3). Mutation of the FACE (Zmpste 24) metalloproteinase, which catalyses the final cleavage step of lamin A processing, results in Restrictive Dermopathy (RD) that also has progeroid features (4). It is the affect of the aberrantly farnesylated prelamin A proteins on nuclear architecture and nuclear function that is presumed to be responsible for the devastating phenotypes of these progeroid diseases (5,6).

At the cellular level, progerin appears to incorporate into the lamina at the nuclear periphery (1,7). In usual cell culture conditions, the resultant nuclei often develop an abnormal and irregular shape with a thickened lamina, especially as the cells reach later passages. There are also various reports of altered distribution of Lamin B (8), nuclear pores, and inner membrane proteins in lamin A mutant cells (3,8).

A loss of peripheral electron-dense heterochromatin is seen by electron microscopy (e.m.) in later passage (p26) HGPS fibroblasts (7). Globally, most HGPS fibroblasts appear to have reduced levels of the heterochromatin protein HP1α and histone modifications (H3K9me3 and H3K27me3) associated with heterochromatic states (8-10). But genomic profiling in HGPS cells suggests a more complex redistribution of H3K27me3 rather than a simple loss (11). H3K9me3 levels in Zmpste 24 deficient cells seems to depend on passage number (12). These chromatin phenotypes are recapitulated in wild-type cells which ectopically express LAΔ50 (7), but they are not rescued in HGPS cells transfected with wild-type lamin A (8). These data serve to underpin the idea that LAΔ50 acts as a dominant negative, or gain of function, mutation. Indeed the presence of progerin may interfere with processing of wild-type lamin A (7).

The presence of progerin, unprocessed prelamin A, or indeed the accumulation of farnesylated prelamin A in *Zmpste24*^-/-^ cells, has been reported to lead to defects in the DNA damage response and in DNA repair and, as a consequence, an increase in DNA damage (13-18). This has lead to the hypothesis that an accumulation of unrepaired DNA damage may be responsible for the premature ageing phenotypes in HGPS, RD and in FACE/Zmpste24 mutants (19,20). This is consistent with the idea that defective DNA repair and increased DNA damage are causally related to both other progeroid syndromes in which known genes of the DNA repair and DNA damage response pathways are mutated (21-23), and also to normal ageing (24,25).

How lamin A mutation and lamin A processing defects might result in defective DNA repair is not clear. Because progerin seems to have an increased association with the nuclear lamina and a decreased association with internal lamin A foci, it could sequester replication proteins away from these internal sites (26) leading to stalled and then collapsed replication forks (24-27). Inappropriate sequestration of repair proteins to these sites of DNA breaks may also be involved (13). Increased generation of reactive oxygen species (ROS) has also been reported in progeria fibroblasts (28). Finally, organisation at the nuclear periphery, especially of heterochromatin, may be important to physically shield the genome from incoming mutgens (29).

There is a clear need to better understand the nature of any defect in DNA repair in HGPS. However, with the exception of some comet assays, previous studies have relied upon markers of the DNA damage response (e.g. γ-H2A.X), rather than DNA damage per se, to address this question. The consequence of unrepaired DNA damage is mutation. Here, using a mutation reporter cell line derived from Muta^TM^Mouse (30), we have directly compared the inherent mutation rate in cells expressing wild-type or LAΔ50 lamin A and also the mutation rate induced in these cells by exposure to exogenous mutagens. We find no significant elevation of mutation rate, scored at the mutation reporter, in LAΔ50/progerin-expressing cells and therefore suggest that underlying the cellular defect in HGPS cells does not lie in defective DNA repair per se.

## Results

### Expression of wild-type and mutant lamin A in mutamouse cells

To investigate direct effects of progerin on the repair of DNA damage we took advantage of a well-established mutation reporter system. In the Muta^TM^Mouse there are ~ 30 copies of λgt10lacZ inserted into the mouse genome (31,32) and *lacZ* then functions as the main target sequence for scoring mutations (33). A stable epithelial cell line, FE1, was established from these animals and is suitable for measurement of endogenous mutation rates, as well as the rates induced in response to a variety of mutagens (30). To characterize the genomic context of the mutation reporter, we used fluorescence in situ hybridization (FISH) with a λgt10lacZ probe on metaphase chromosomes from FE1 cells. In each spread three chromosomes were labeled by the probe, and analysis of their DAPI-banding pattern suggested that these might be *Mus musculus* chromosomes 3 (MMU3). Combined analysis with a chromosome paint for MMU3 confirmed this (Fig. 1A). We conclude that in the aneuploid FE1 cell line (n=78 chromosomes/spread) there has been a triplication of the original λgt10lacZ containing chromosome.

**Figure 1.**
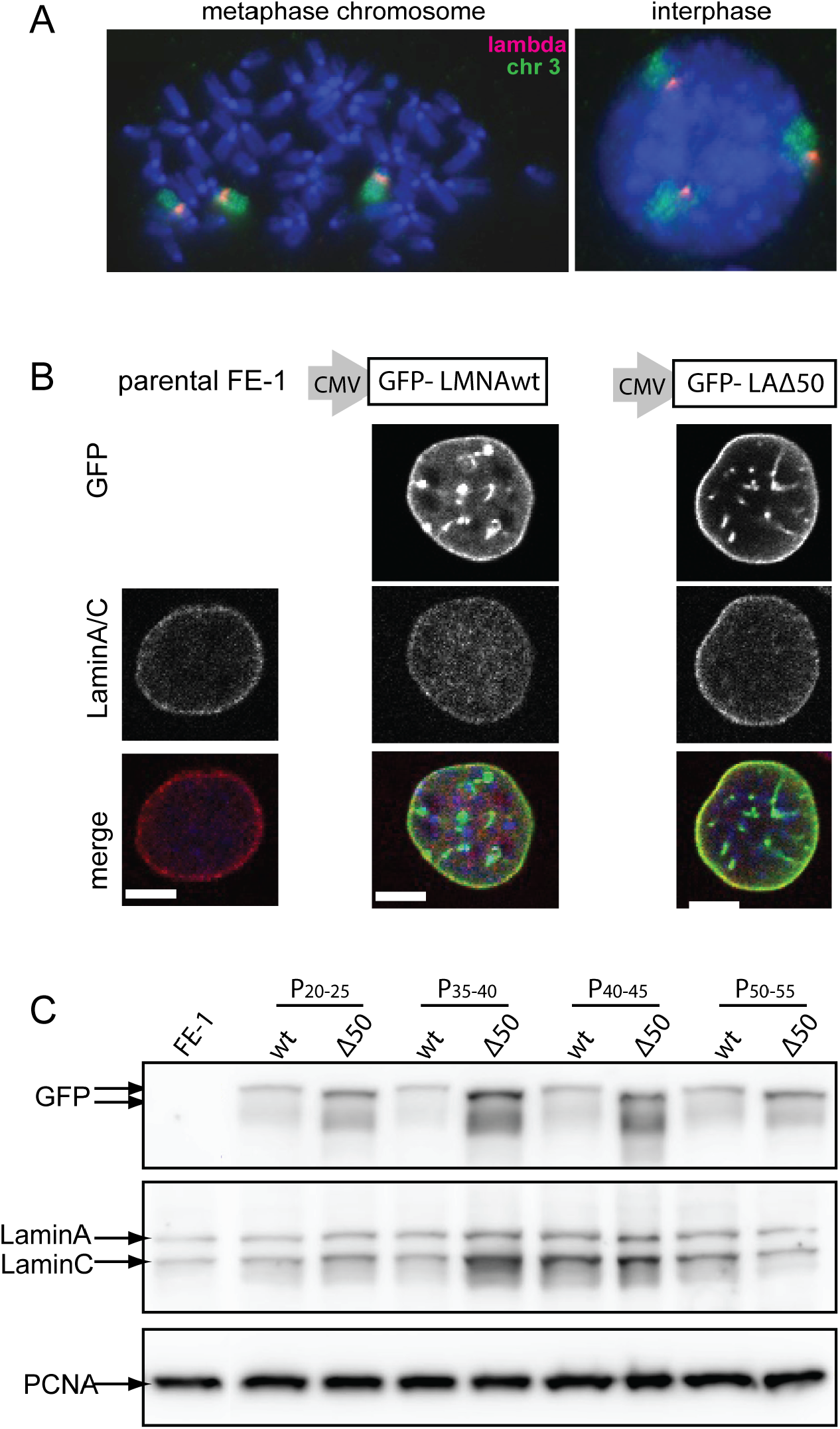
Expression of mutant and wild-type lamin As in mutamouse cells A) FISH on metaphase chromosome spread (left) or interphase nucleus (right) from FE1 Mutamouse cells hybridized with probes for phage X DNA (red) and mouse chromosome 3 (green). DNA is counterstained with DAPI. B) Immunofluorescence on parental FE1 cells (left column) and FE1 stable transfectants expressing wild-type (middle column) or A50 (right) mutant lamin Afused to GFP. Confocal mid planes showing - top row: GFP fluorescence, middle row: immunofluorescence for laminA/C, bottom row: merge (GFP/ green, laminA/red and DAPI/blue). C) Western blöt of protein extracts from parental FE1 cells (left) and FE1 stable transfectants expressing wild-type (wt) or A50 mutant lamin Afused to GFP, and harvested at various passages. Membranes were probed with antibodies recognising; (top row) GFP, (middle) lamin A and lamin C, (bottom) PCNA-as a loading control.

To determine the effects of progerin expression on mutation rate, we created FE1 cell lines stably expressing GFP-tagged human lamin A – either wild-type (wt), or ΔLA50/progerin (7). Ectopic expression of N-terminally GFP-tagged lamin A has been shown to result in the stable integration of the tagged protein into the lamina at the nuclear periphery (34). To ensure stable expression from the transgene, we FACs-sorted GFP-expressing cells. Fluorescence microscopy confirmed that both the wt and ΔLA50 mutant GFP-tagged lamin As concentrate at the nuclear lamina. Optical sectioning along the *z* axis showed that any bright, apparently internal, foci of GFP-lamin A were due to invaginations of the nuclear periphery (Fig. 1B).

Immuno-blotting confirmed the stable expression of the GFP-tagged lamin A over prolonged time in cell culture, with no accompanying obvious decrease in expression of endogenous lamin A (Fig. 1C).

### Heterochromatin and nuclear morphology in lamin A expressing mutamouse cells

Reduced levels of heterochromatic histone modifications, especially H3K9me3, and the heterochromatin protein 1 (HP1α) that binds to this mark, have been reported in HGPS cells (8) and in human cells ectopically expressing ΔLA50 (9). By immunoblotting we saw a small reduction of H3K9me3 and HP1α levels in late passage FE1 cells expressing ΔLA50 as compared to cells expressing wild-type lamin A (Fig. 2A), though we did could not detect loss of H3K27me3 in the presence of ΔLA50.

**Figure 2.**
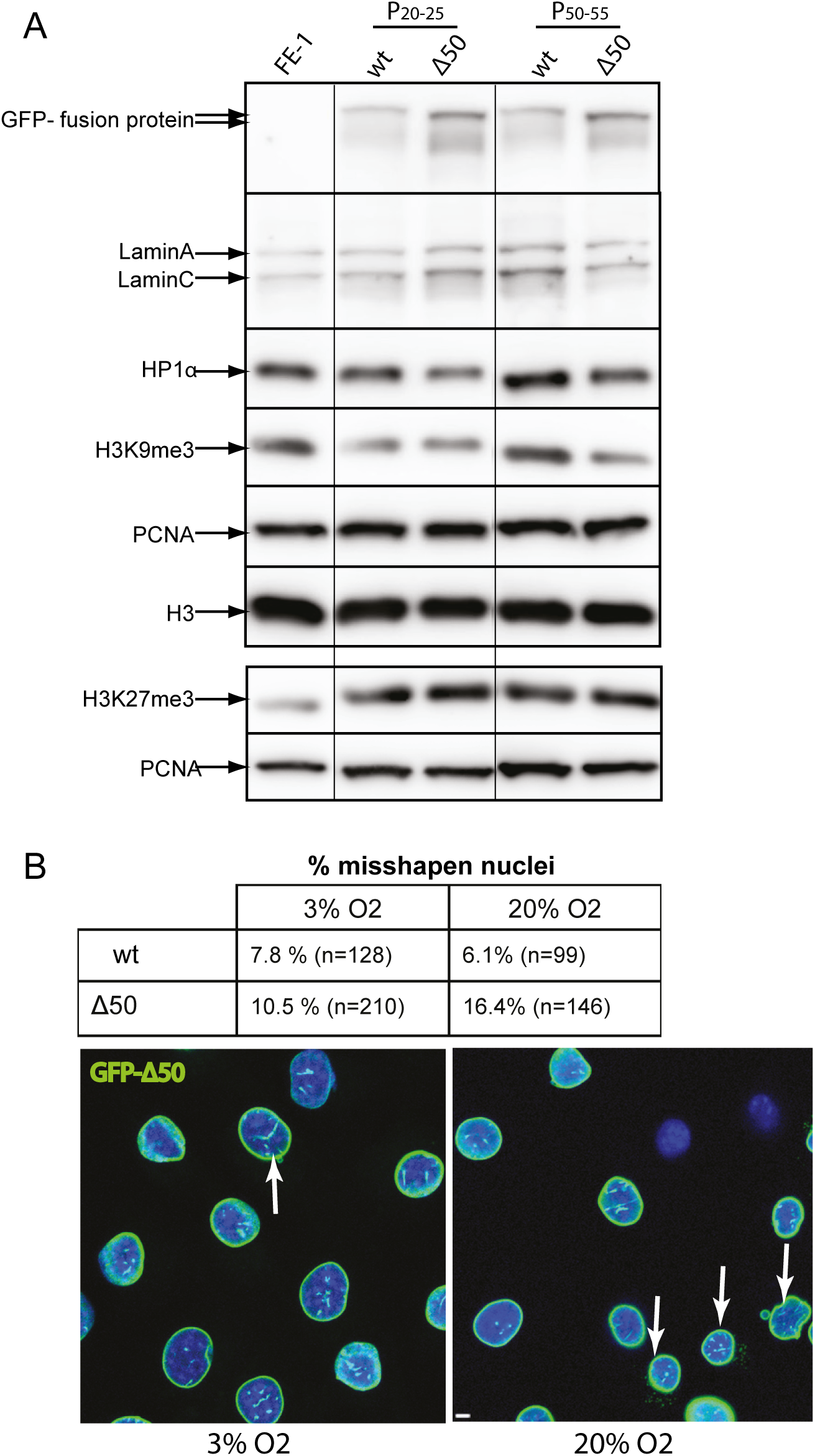
Histone modifications and nuclear morphology in lamin Aexpressing cells A) Immunoblotting of proteins from FE-1 parental cells, and from early (P20-25) and late (P50-55) passage FE-1 stable transfectants expressing wild-type (wt) or A50 mutant lamin A, with antibodies detecting; laminA/C, HP1a, H3K9me3 and H3K27me3. H3 and PCNA serve as loading controls. B) Table shows the proportion of abnormal nuclei scored in FE-1 transfectants expressing wild-type (wt) or A50 mutant GFP-tagged lamin A grown under conditions of low (3%) or high (20%) 02. n= number of nuclei scored. Below, confocal mid planes show examples of fields of A50-expressing cells grown under conditions of low (3%) or high (20%) 02. Green = GFP, blue = DAPI. Arrows indicate nuclei scored as abnormal.

Another characteristic of HGPS is the acquisition of aberrant nuclear morphology in late passage cells. Indeed, under normal cell culture conditions in late passage ΔLA50-expressing FE-1 cells we observed that the proportion of cells with aberrant nuclear morphology (evidence of nuclear blebbing, nuclear fragmentation) was > 2.5 fold higher (16.4%) than in FE-1 cells expressing wild-type lamin A (6.1%) (Fig. 2B). Thus we conclude that FE-1 transfectants exhibit many of the gross nuclear features reported for HGPS cells and other cells transfected with progerin-expressing constructs and that they therefore represent a suitable model system for studying the effects of ΔLA50 on DNA damage and mutation.

It is known that mouse cells grown in standard tissue culture conditions – i.e. under high oxygen tension (20% O_2_), are subject to oxidative stress and have an elevated mutation rate (35). Since we wanted to study the effects of ΔLA50 on intrinsic and induced mutation rates we did not want to work against a background of mutations introduced as a consequence of non-physiological culture conditions. Therefore, once established, we maintained the FE-1 cells and the WT and ΔLA50-expressing FE-1 lines under physiological O_2_ (3%) for multiple passages. The mutation rate of mouse cell lines grown under these conditions, is reported to be the same as for primary cells (35).

Under these conditions, we noted that the proportion of ΔLA50 expressing cells exhibiting aberrant nuclear morphology was markedly reduced (10.5%) relative to the same cells grown in 20% O_2_ and was only slightly greater than cells expressing wild-type lamin A (Fig. 2B). Thus it appears that the aberrant nuclear morphology of lamin A mutant cells may be due to their increased sensitivity to the cell stress and DNA damaging effects of abnormally high O_2_ concentration. This would be consistent with the increased sensitivity to ROS reported for HGPS fibroblasts (28).

### Intrinsic mutation rate in the presence of wild-type and mutant laminA

We first assessed the intrinsic spontaneous mutation rate in lamin A FE1 transfectants by recovery and packaging of λgt10-lacZ genomes from the cellular DNAs of parental FE-1 cells, and cells stably expressing wild-type or Δ50 lamin A. Resultant phage containing *lacZ* mutations were selected for by infection of galE- *E.coli* (BIK12001) and plating on minimal agar containing 0.3% w/v phenyl-*β*-D-galactosidase (P-Gal) (36,37). In wild-type (lacZ^+^) phage, release of the galactose moiety from P-Gal by β-galactosidase results in the accumulation of toxic UDP-galatose in GalE- strains. Therefore only cells infected by lacZ-mutant phage survive and form plaques (38) (Fig. 3A). Mutation frequency is then expressed as the ratio of mutant plaques (+P-Gal plates) to total plaque forming units (pfu) on non-selective plates. The efficacy of selection was first tested using known wild-type and (L1A15) mutant stocks of λgt10-lacZ phage (37). There was a 10^4^ fold drop in plating efficiency on PGal selective plates for the wild-type over lacZ- mutant phage, comparable to previous reports using this system (37) (Fig. 3B and C).

**Figure 3.**
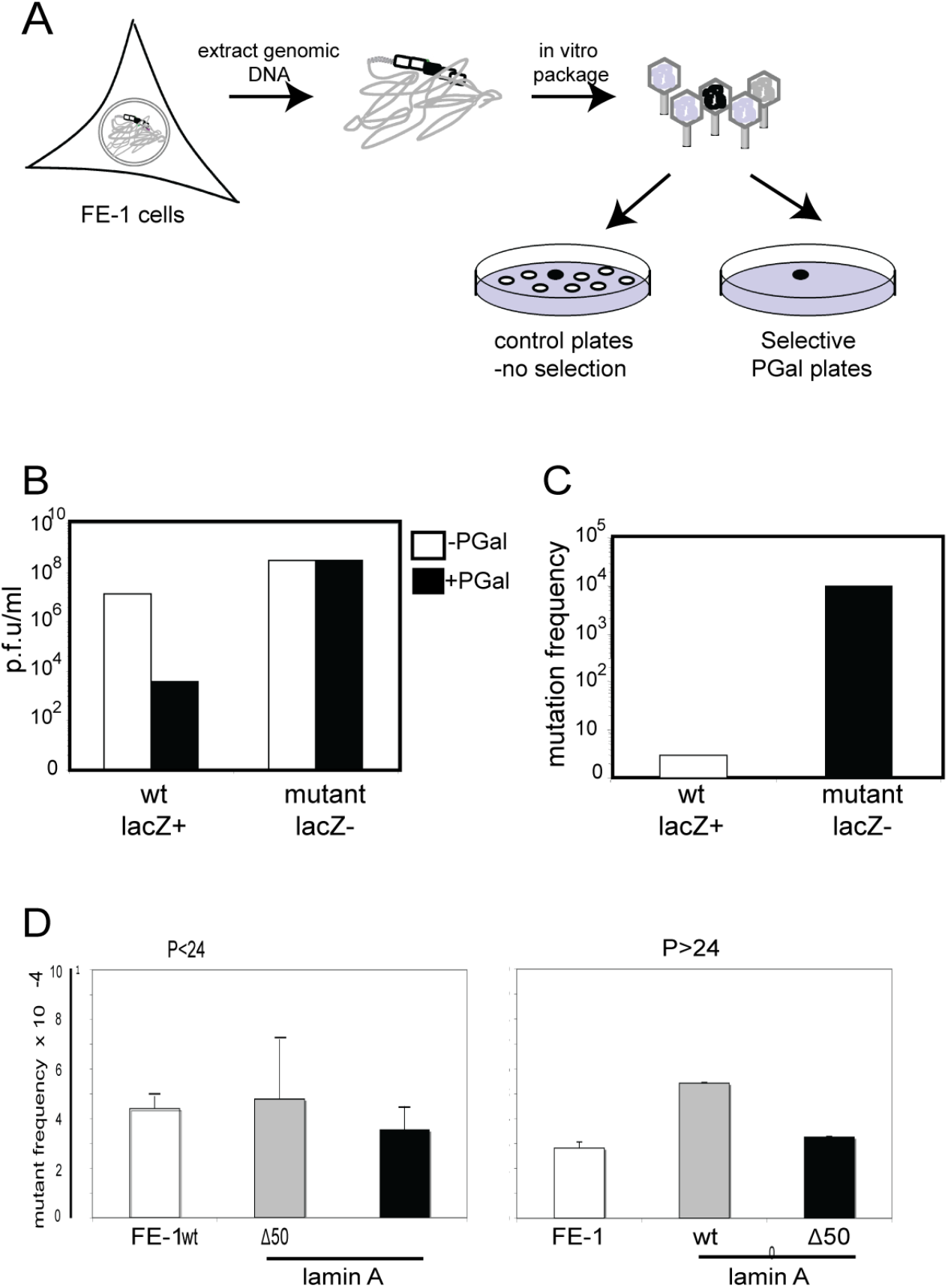
Determination of intrinsic mutant frequency in lamin A expressing cells A) Schematic showing the determination of mutation frequency atXgtlOlacZ sequences in FE-1 cells, by in vitro packaging of phage DNA and plating of infected E.coli on PGal selective plates. Only phage with mutations in lacZ (black filled) grow on selective PGal plates. Mutation frequency = ratio of pfu on PGal/pfu on non-selective (-Pgal) plates. B and C) Confirmation of efficacy of PGal selection. B) Graph shows plating efficiency (pfu/ml in log scale) of wild-type (lacZ+) and known mutant (LacZ-) phage stocks on selective (+PGal, black bars) and non-selective (-PGal, white bars) plates. C) Mutant frequency (pfu +PGal/pfu -Pgal) measured for wild-type (white bars) and known mutant (LacZ-) stocks of phage. D) Mutant frequency (x104) measured at the Xgt1 OlacZ transgenes in FE-1 cells and in these cells stably expressing wild-type lamin A (wt) and HGPS mutant lamin A (A50). Cells were tested at low passage (< P24, left hand graph) and then again at higher passage number (P=48-53). Graphs show the mean (± s.e.m) for genomic DNAs isolated from two independent experiments, and with technical replicates for packaging of these DNAs.

Mutant frequency from parental FE-1 cells, and lamin A transfectants was then assessed at early (<passage 24) and later (P48-53) passages. The intrinsic mutant frequency of all three cell lines was low (<5.5 x 10-4) and there was no evidence for an increase in transfectants stably expressing mutant lamin A (Δ50) (p=0.67) (Fig. 3D). Indeed, there was some evidence that the mutant rate in cells expressing wild-type lamin A was slightly elevated relative to both the parental FE-1 cells and the Δ50 cell line, but this was not statistically significant.

### Induced mutant rate in the presence of wild-type and mutant lamin A

To determine if cells expressing mutant lamin A were more susceptible to DNA damage induced by exogenous mutagens, we exposed FE-1 and lamin A transfectants to two different mutagens.

We first analysed the effects of exciting, but non-ionising, UV-C (254nm) radiation (39). This induces cyclobutane pyrimidine dimers (CPDs) and strand breaks (40,41). Cells were exposed to 10 J/m2 UV 254 nm on two consecutive days and harvested 72h after the first exposure to mutagen. Efficacy of the mutagen was assessed by measuring the mutant frequency of phage particles recovered from parental FE-1 cells before, and after, treatment. UV-C elevated the mutant frequency 10 fold (p < 1 x 10-6)(Fig 4A). UV-C similarly elevated the mutant phage frequency from FE-1 transfectants expressing wild-type or mutant (Δ50) lamin A. However, as seen for the intrinsic mutation frequencies, and contrary to what was expected from proposed models of HGPS pathogenesis, the mutant frequency in cells expressing wild-type LMNA appeared a bit higher than in cells expressing LAΔ50 mutant protein, although overall this was not statistically significant p = 0.14 (Fig. 4B).

**Figure 4.**
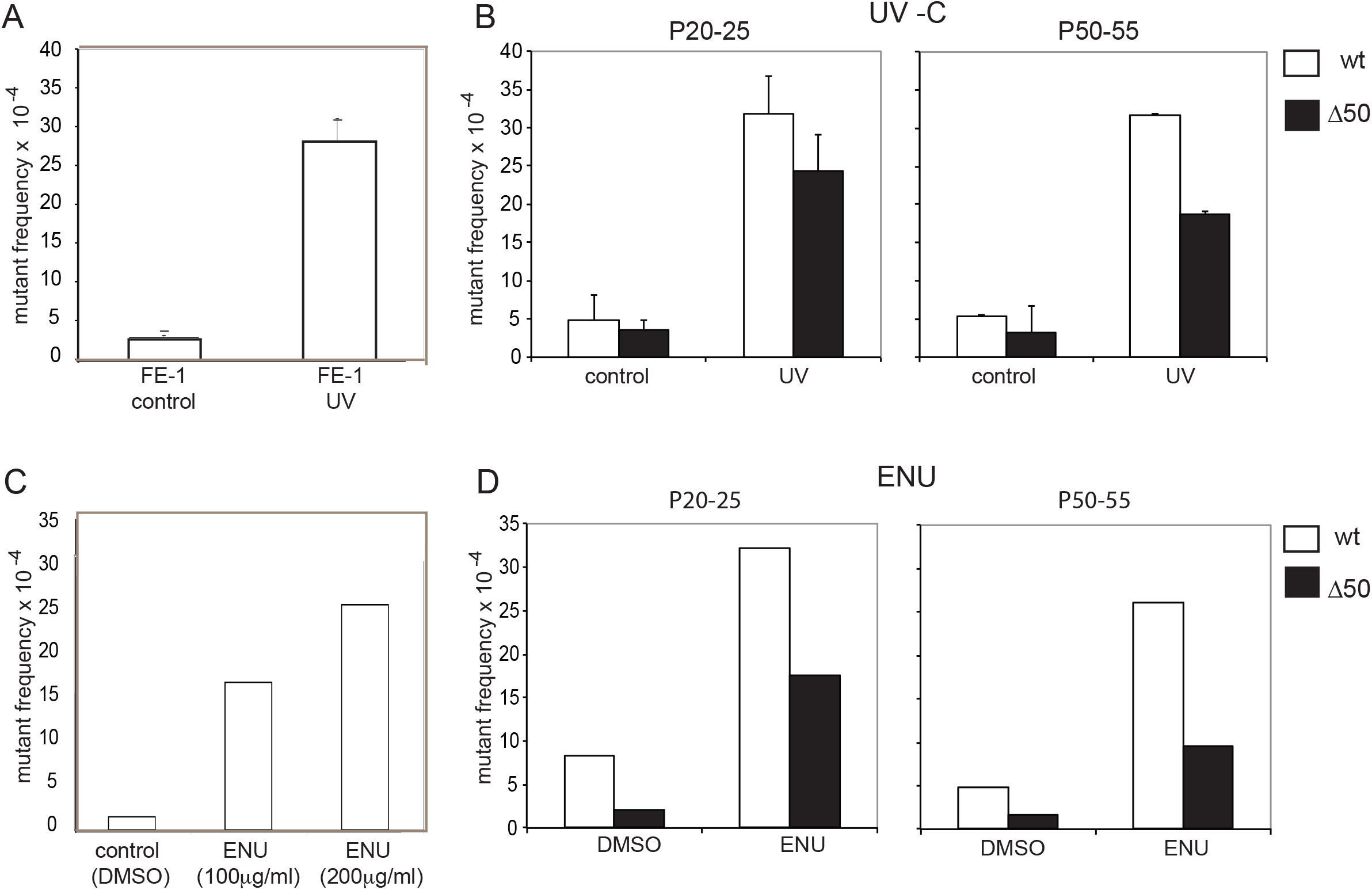
Induced mutant frequency in wild-type and mutant lamin A expressing cells A) Mutant frequency (x104) measured at the Ågt10lacZ transgenes in untreated control FE-1 cells and in cells exposed to 10 J/m2 UV 254nm. B) Mutant frequency (x104) in control and UV treated FE-1 cells stably expressing wild-type lamin A (wt, white bars) or Δ50 mutant lamin A (black bars). Cells were tested at low passage (< P24, left hand graph) and then again at higher passage number (P=48-53). Graphs show the mean ± s.e.m for genomic DNAs isolated from two independent experiments, and with technical replicates for packaging of these DNAs. C) Mutant frequency (x104) measured at the Ågt10lacZ transgenes in control (DMSO) FE-1 cells and in cells exposed to 0.85 and 1.7mM (100 and 200 µg/ml) ENU. D) As in B but for control (DMSO) and 1.7mM ENU treated FE-1 cells stably expressing wt or A50 mutant lamin A.

To ascertain whether a similar result could be obtained using a different mutagen with a different mode of action, we treated cells with the alkylating agent ethyl nitrosourea (ENU). ENU induces mainly A:T to T:A transversions, probably due to thymine adducts. We treated FE-1 cells with 0.85 and 1.7mM (200µg/ml in DMSO) ENU, since the latter is the dose reported to produce a maximal mutant rate of 2 x 10-3, i.e. 5 x spontaneous rate, at the lacZ reporter from MutaTMmouse (27). Indeed, we saw a similar increase in mutant frequency, compared to DMSO treated controls, in our assay (Fig. 4C).

The mutant frequency in lamin A-expressing FE-1 stable transfectants was also elevated by ENU treatment but, as was seen for UV treatment, the ENU-induced mutant rate was higher in the cells expressing wild-type lamin A than those expressing LAΔ50 and this was statistically significant (p = < 1 x 10-4)(Fig. 5D). Hence, by scoring for mutant lacZ in transgenic murine cells, we are unable to find any evidence for an increased rate of unrepaired DNA damage caused by the presence of progerin. If anything the expression of progerin suppressed the mutation frequency, at least in the case of the mutagen ENU.

**Figure 5.**
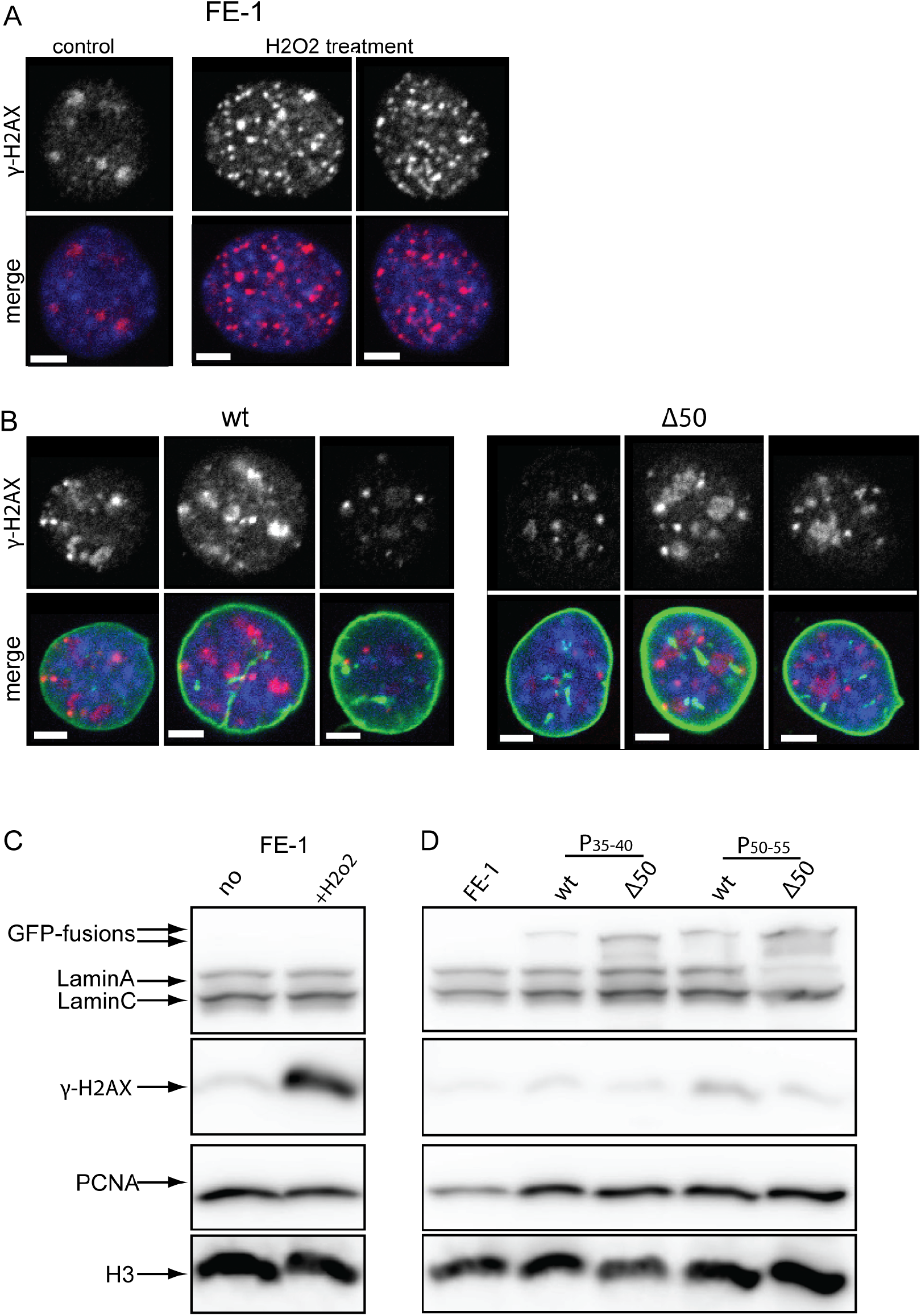
Indicators of the DNAdamage response in wild-type and mutant lamin Aexpressing cells A) Immunofluorescence with antibody detecting yH2A.X in FE-1 cells and cells exposed to 1 mM H202 for 25min. In merge antibody is red, blue = DAPI. B) Immunofluorescence with antibody detecting yH2A.X in FE-1 cells expressing wild-type (wt) or A50 mutant lamin A. In merge antibody is red, green = GFP, blue = DAPI. C) Immunoblot with antibodies detecting lamin A/C and yH2A.X in FE-1 cells and cells exposed to 1mM H202 for25min. PCNAand H3 serve as loading controls. D) Immunoblot with antibodies detecting lamin A/C and yH2A.X in FE-1 cells and transfectants expressing wild-type (wt) or A50 mutant lamin A.

### No increase in γH2A.X in progerin-expressing cells

An increased appearance of γH2A.X and 53bp1 foci, and defective recruitment of 53bp1 to sites of DNA damage, has been reported in Zmpste24 deficient mouse embryonic fibroblasts (MEFs), in HGPS fibroblasts, and in wild-type MEFs expressing unprocessed prelamin A (13,14), suggesting a perturbed response to DNA damage and compromised DNA repair. To investigate this further we first showed that the appearance of γH2A.X foci could indeed be induced by exposure of FE-1 cells to oxidative damage (H_2_O_2_) and detected both by immunofluorescence (Fig. 5A) and immunoblotting (Fig. 5C). However, under low O2 growth conditions we did not find evidence, by either immunofluorescence (Fig 5B) or by immunoblotting (Fig. 5D), for elevated levels of γH2A.X in progerin expressing FE-1 cells. Some increase in γH2A.X was seen with prolonged passage in culture, but this was more pronounced in cells expressing wild-type lamin A than in those expressing the mutant protein.

## Discussion

There is little doubt that the accumulation of damage to cellular components, including DNA, contributes to normal chronological aging (42). Both genome instability and point mutations increase with age (43). The premature aging phenotypes seen in many genetic disorders with mutation of genes involved in DNA metabolism reaffirms that DNA damage and a loss of genome integrity is a key feature of both normal and abnormal aging (24).

The mechanisms that lead to a premature aging phenotype due to the dominant LAΔ50 mutation in HGPS are a matter of ongoing debate. However, one plausible mechanism is that there are defects in DNA damage sensing and/or DNA repair (44). Studies that have addressed this possible mechanism have examined markers of the DNA damage response and DNA repair pathways. However, compromised sensing or repair of DNA damage should lead to an increase in un-repaired DNA damage – i.e. mutation. This has been inferred (13) but not previously directly assessed, in HGPS cells or in cells expressing progerin (LAΔ50). Here we have used transgenic reporter cell lines to directly investigate the affects of LAΔ50 mutant lamin A on the spontaneous and induced mutation rate of the phage mutation reporter integrated into the genome of these cell lines (Fig 1). Such transgenic mutation reporters have been used to reveal the accumulation of mutation load during chronological aging (45-47), but to our knowledge have not previously been used to assess the role of nuclear structure on DNA damage.

We found no evidence for an increase in the spontaneous mutation rate (Fig. 3) in cells expressing human progerin mutant lamin A compared with cells expressing wild-type lamin A. Many different DNA repair pathways are mutated in other progeria syndromes and all DNA repair pathways seem to decline with chronological aging (43). The mutagens that we have investigated here (UV and ENU) likely are targeted by global genome nucleotide excision repair (NER), base excision repair (BER) and break repair. NER is a ’detect-excise-and-patch’ repair system for a broad class of helix-distorting lesions such as UV-induced photoproducts and many bulky chemical adducts. The global-genome NER (GG-NER) pathway detects such lesions anywhere in the genome, whereas the transcription-coupled NER (TC-NER) pathway is selective for lesions that block transcriptional elongation. UV-induced CPD photoproducts removal is reduced in older and senescent fibroblasts (43), especially in non-transcribed region (44), and BER activity is reported to be decreased in old mice (45). Similarly MEFS from lamina-associated progeroid syndromes are reported to be hypersensitive to UV (14). We did score a significantly elevated mutant rate at FE-1 cells and transfectants after their exposure to either UV or ENU, however this induced mutant rate was no higher in FE-1 transfectants expressing LAΔ50 than in the parental cells or cells expressing wt lamin A (Fig. 4).

Because we have only scored mutation at an unexpressed reporter locus (LacZ), we cannot exclude that it is specifically transcription-coupled repair that is affected in HGPS, though no increase in spontaneous mutation rate has been seen in progeroid TC-NER mutants (51). Similarly, since not all tissues seem to age prematurely in HGPS it is possible that the FE-1 cell line is not from a suitable tissue-type to reveal a defect in DNA repair caused by expression of LAΔ50 (52).

As has been previously reported (3,7,9), we found a increased proportion of abnormally and irregularly shaped nuclei in cells expressing LAΔ50 mutant as compared to wt lamin A (Fig. 2). However, we noticed a decrease in this proportion when LAΔ50 expressing cells were grown at physiological oxygen levels (3%) as compared to the 20% O2 of standard tissue culture. A similar decrease in aberrant nuclear morphology was not seen in cells expressing wt lamin A. This suggests that progerin-expressing cells may be especially sensitive to oxidative stress and this is consistent with the elevated levels of reactive oxygen species reported in HGPS cells (28, 53).

## Materials and methods

FE1 Muta^TM^Mouse lung epithelial cells were cultured as described by White et al. (30). Briefly, cells were cultured in 1:1 DMEM:F12 nutrient mixture (Invitrogen) supplemented with 2% FBS (Sigma), 2 mM glutamine, and 1 ng/ml murine epidermal growth factor (Invitrogen). Cells were maintained at 37^o^C in hypoxic incubators (New Brunswick Galaxy 170R) with 3% O_2_ or in atmospheric O_2_ levels incubators when indicated. For hydroxygen peroxide (H_2_O_2_) treatment, cells were exposed to 1mM H_2_O_2_ for 25min.

### FISH

Cells were swollen in 0.56% KCl before fixation in 3:1 methanol:acetic acid. Slides were incubated with 100ug/ml RNaseA in 2 x SSC for 1 hr, washed in 2 x SSC and dehydrated through an alcohol series. Slides were then denatured in 70% formamide/2xSSC for 1min. λ DNA was purified using manufacturer’s protocol (Qiagen, # 12543) and labelled by nick translation with digoxigenin-11-dUTP (54). Approximately 100 ng of labelled DNA probe and 15 µl of FITC-mouse chromosome-3 paint (Cambio) were used per slide, together with 5 µg of mouse Cot1 DNA (GIBCO BRL) and 5 µg salmon sperm DNA. Probes were denatured at 70°C for 5 min, reannealed with Cot1 DNA for 15 min at 37°C and hybridized to the denatured slides overnight. Digoxigenin-labelled probes were detected with Texas-Red anti-sheep (Vector). Slides were counterstained with 0.5 µg/ml 4’,6’-diamidino-2-phenylindole DAPI in Vectashield.

### Cloning and expression of lamin A and LAD50

Stable FE-1 cells were prepared expressing either EGFP-myc-LA or EGFP-myc-hLAΔ50. The coding regions from pTRE-EGFP-myc-hLMNA or –hLMNAΔ50 (7,9) were excised as a 2.8 kb BamHI/XbaI fragment and cloned into pcDNA3.1 (+) (Invitrogen). The resulting plasmids (pcDNA-EGFP-myc-hLMNA and pcDNA-EGFP-myc-hLMNAΔ50) were linearised with ScaI and transfected into FE-1 cells by lipofectamine 2000 (Invitrogen). After 48 hours, cells were trypsinised and GFP-expressing cells sorted using a BD FACSAriaII SORP instrument (Becton Dickinson). The 488nm laser was used for measuring forward scatter, side scatter and GFP fluorescence (525/50nm bandpass filter). BD FACSDiva software (Becton Dickinson, Version 6.1.2) was used for instrument control and data analysis. The purified GFP-expressing cells were seeded in 6-well plate and expanded. Cell populations were sorted regularly for GFP fluorescence to maintain a high-proportion of GFP-lamin A expressing cells. After establishment of stably expressing cell lines, cells were maintained in a 3% O2 incubator.

### Immunofluorescence

Cells grown on coverslips were fixed in 4% paraformaldehyde in PBS and permeabilised in 0.2% Triton X-100/PBS for 12 mins. Fixed cells were incubated overnight at 4^o^C with primary antibodies against Lamin A/C (1:100, Cell Signaling) or γ-H2AX phospho-[Ser139] (1:100, #05-636 Millipore). The slides were then incubated for 1 hour at room temperature with Alexa-Fluor 488 or Alexa-Fluor 594 secondary antibodies (1:1000, Invitrogen). Cells were counterstained with 0.02 µg/ml DAPI in PBS and mounted in Vectashield. Slides were examined on a Nikon A1R confocal microscope equipped with a CFI Plan Fluor 40x/1.30 oil objective and NIS elements software.

### Western blotting

Western blot analysis was carried out using standard protocols. Immunoblotting was performed with primary antibodies directed against Lamin A/C (1:2000, #sc-6215 Santa-Cruz Biotechnology) or (1:1000, #4777, Cell Signaling), GFP (1:2000, #11-814-460-001 Roche), H3K27me3 (1:2000, #07-449 Millipore), H3K9me3 (1:1000, #05-1242 Millipore), γ-H2AX phospho-[Ser139] (1:500, #05-636 Millipore), Hp1α (1:500, #05-689 Millipore), PCNA (1:20000, #sc-56 Santa-Cruz Biotchnologies) and H3 (1:50000, Millipore). Blots were detected by horseradish peroxidase (HRP)-conjugated donkey anti-rabbit, anti-mouse or anti-goat whole molecule IgG (1:10000) and chemiluminescence using ChemiGlow West (Alpha Innotech). Signal was analysed using ImageQuantTL LAS4010 (Version 1; GE Healthcare).

### Measurement of LacZ mutant frequency

3x10^5^ cells were cultured in a 100 mm culture dish overnight. For UV-C treatment, cells were washed with PBS and exposed to 10 J/m2 (UV Stratalinker 1800, Stratagene) at 254 nm, which was repeated the following day. Cells were collected 72h after the first exposure to mutagen. For N-ethyl-N-nitrosourea (ENU, Sigma) treatment, cells were incubated for 6h in serum-free medium containing 200µg/ml ENU diluted in DMSO, washed with PBS and then incubated for 72h with complete medium. Cells were lysed for 3h at 55^o^C in 100 mM Tris pH 8, 200 mM NaCl, 5mM EDTA, 0.2% SDS and 0.1 mg/ml proteinase K. DNA was extracted with phenol/chloroform (1:1), followed by chloroform and precipitated by isopropanol, and resuspended in 25 µl of TE.

λgt10lacZ DNA were rescued from FE1 genomic DNA using the Transpack^TM^ lambda packaging system (Stratagene) according manufacturer’s instructions. The phage preparation was used to infect E. coli galE^¯^ cells prepared in 10mM MgSO4 at an OD600nm = 2. The LacZ mutant frequency was determined by using the P-Gal-positive selection assay (36). Briefly, 100 µl packaged phage particles were incubated with 0.5 ml of host bacterium for 20 min at 37^o^C and plated on agar containing 0.3% w/v Phenyl β-D-galactoside (P-gal, Sigma, # P6501). After incubation overnight at 37^o^C, the number of LacZ mutant plaque-forming units (pfu) was counted. Concurrent pfu titers on non-selective agar were employed to calculate total pfu. Mutant frequency is expressed as the ratio of (pfu of PGal: total pfu (no PGal). The statistical significance of differences in mutant frequency scored between cell lines was assessed using t-tests.

## Acknowledgements

This work was supported by an ERC advanced grant (no. 249956) and by the Medical Research Council, UK. We thank Elisabeth Freyer (MRC HGU technical services) for FACs sorting. We are grateful to Bob Goldman (Northwestern University, Chicago) for the gift of lamin A constructs and to Ichizo Kobayashi and Naofumi Handa (University of Tokyo) for wild-type and mutant λgt10lacZ phage stocks. The FE1 cell line from Muta^TM^mouse and GalE- E. coli were provided by Health Canada, Ottawa (Paul White and George Douglas). We are also grateful to Paul White (Health Canada) for helpful comments on the manuscript.

